# Population density modeling of mixed polymorphic phenotypes: an application of spatial mark-resight models

**DOI:** 10.1101/2020.06.08.129197

**Authors:** Abishek Harihar, Dipankar Lahkar, Aparajita Singh, Sunit Kumar Das, M Firoz Ahmed, Ramie H Begum

## Abstract

Melanism is a form of pigmentation polymorphism where individuals have darker colouration than what is considered the “wild” phenotype. In the case of leopards, *Panthera pardus*, melanism occurs at higher frequencies amongst populations in tropical and subtropical moist forests of south and southeast Asia, presenting a unique challenge in estimating and monitoring these populations. Unlike the wild phenotype that is readily recognizable by its rosette patterns, melanism results in individuals being unidentifiable or ‘unmarked’ through photographic captures obtained using white flash cameras. Spatial mark-resight (SMR) models that require only a subset of the population to be ‘marked’ offer the opportunity to estimate population density. In this study, we present an application of SMR models to estimate leopard densities using camera trap survey data from three sampling years at Manas National Park (MNP), India. By using an SMR model that allowed us to include captures of unidentified sightings of marked individuals, we were also able to incorporate captures where identity was either not confirmed or only known from a single flank. Following 18,674 trap-days of sampling across three years, we obtained 728 leopard photo-captures, of which 22.6% (165) were melanistic. We estimated leopard densities of 4.33, 2.61 and 3.37 individuals/100km^2^ across the three years. To our best knowledge, these represent the first known estimates of leopard densities from populations comprising both melanistic and wild phenotypes. Finally, we highlight that SMR models present an opportunity to revisit past camera trap survey data for leopards and other species such as Jaguars, *P. onca*, that exhibit phenotypic polymorphism towards generating valuable information on populations.

## Introduction

Melanism is a form of pigmentation polymorphism where individuals have darker colouration than what is considered the natural or “wild” phenotype (Majerus 1998). Amongst Felids, melanism is believed to have evolved independently several times, being documented in 14 of 40 species, and is likely associated with several ecological, physiological and behavioural factors (Eizirik et al. 2003; Schneider et al. 2012, 2015; Graipel et al. 2019). In the case of leopards, *Panthera pardus*, melanism is induced by a recessively inherited mutation in the ASIP (agouti signalling protein) gene and occurs at a range-wide frequency of ~11% (Schneider et al. 2012; da Silva et al. 2017). Although melanistic individuals most often occur as a polymorphic phenotype, some populations exhibit near fixation of this recessive allele (Kawanishi et al. 2010). As Tropical and Subtropical Moist Broadleaf Forests of south and southeast Asia harbour higher frequencies of melanism (da Silva et al. 2017), this presents a unique challenge in estimating and monitoring populations in these regions.

As the “wild” phenotype is individually identifiable by rosette patterns, photographic capture-recapture sampling is regarded as the most robust and cost-effective method of estimating population size and density of leopards (Balme et al. 2009). However, the masking of rosettes, caused by melanism (Schneider et al. 2012), results in a proportion of individuals remaining ‘unmarked’. Hedges et al., (2015) discovered that by forcing infrared flash camera traps (IR camera traps) into night mode, the rosettes of melanistic leopards were revealed allowing for individuals to be identified and population density estimated using spatial capture-recapture (SECR) analysis. While the use of IR camera traps may be inevitable at sites where a near fixation of melanism has been observed (Kawanishi et al. 2010), white flash cameras that produce high-quality images (with no motion blur) are considered ideal for capture-recapture studies as they allow for more reliable identification of individuals from coat patterns (Rovero et al. 2013). Therefore, at sites where both phenotypes occur, mark-resight models that require only a subset of the population to be ‘marked’ present an opportunity in estimating population size and density with white flash cameras.

Mark-resight methods have been in use for some time (McClintock & White 2012), and more recently spatial mark-resight (SMR) models have also been developed (Chandler & Royle 2013; Sollmann et al. 2013; Efford & Hunter 2018). These models estimate population density under the assumption that both marked and unmarked individuals have the same probability of being encountered in the re-sighting occasions. A rare circumstance when this assumption is satisfied is when some individuals carry natural marks, and others do not (Rich et al. 2014). However, unlike in studies where the number of individuals that are physically marked and released before the resighting session is known, in cases where individuals are identified by natural markings (as with leopards), the number of marked individuals is unknown and needs to be estimated from resighting data (Royle et al. 2014; Efford & Hunter 2018). Another assumption is that marked individuals are always correctly identified. Often during camera trap surveys, partial identities are obtained (i.e. right-only and left-only flank captures) and/or marks are unreadable (owing to motion blur obscuring marks or marks being hidden from view; Figure A1). Although the inclusion of this information can have subtle effects on the estimate of density, recent developments in SMR models allow for the incorporation of such data in the estimation process (e.g. Efford and Hunter, (2018)), ensuring no loss of capture data.

In this study, we present an application of SMR models to estimate leopard densities at Manas National Park (MNP), India, where the population comprises of both wild and melanistic phenotypes (Figure 1). Furthermore, as we use data from three sampling years, we explore the effects of pooling parameter values that could be common across years in a multi-session analysis towards improving the precision of our density estimates.

## Methods

### Study area

MNP is an 850 km^2^ protected area along the Indo-Bhutan international border (see Lahkar et al., (2020) & (2018) for further details). It lies within the Tropical and Subtropical Moist Broadleaf Forest biome (Olson et al. 2001), where the observed frequency of melanism is five-fold higher than the expected number based on the global average (da Silva et al. 2017). Since the end of ethnopolitical conflict in the region in 2003, MNP has received much biodiversity research and conservation attention (Borah et al. 2014; Lahkar et al. 2018, 2020). Nevertheless, the population status of leopards is poorly understood. Borah et al., (2014) estimated a density of 3.40±SE 0.82 leopards per 100 km^2^; however, this was based on an SECR analysis of rosetted individuals (excluding melanistic captures).

### Field methods

Leopard photo-capture data for this study were obtained from systematic camera trapping surveys conducted within 500 km^2^ of MNP. Sampling was carried out across three successive cool, dry seasons (December to April) of 2016-17, 2018 and 2018-19. We placed a pair of digital white flash passive camera traps (Panthera, New York, USA, Models V4, V5 and V6) at trap densities of one per 4 km^2^ (in 2016-17 & 2018-19) and 2 km^2^ (in 2018) (Table 1; Figure A1). The cameras were operational for 24 hours a day (Jhala et al. 2020; Lahkar et al. 2020). Our sampling design ensured that the trap array was large so that most individuals were exposed to capture within the trapping grid and trap spacing was small relative to individual space use to accurately estimate detection parameters (Sollmann et al. 2012; Sun et al. 2014). Trap locations were chosen following reconnaissance surveys conducted along forest paths, roads, ridges, dry river beds and animal trails prior to each year’s survey. At each trap location, we placed a pair of cameras (without baits), facing each other. Sampling was restricted to a period of 60 days to ensure demographic closure. We downloaded the photographs from all trap stations and catalogued them using Camera Trap File Manager software (Olliff et al. 2014), at regular intervals (usually twice a week). Finally, we created year-specific databases of leopard captures that specified the date, time and location of capture for further analyses.

### Spatial Mark-Resight analysis

To estimate density, SMR models require two key datasets: the spatial detection histories of ‘marked’ and occasion-specific spatial counts of ‘unmarked’ individuals. To generate detection histories of marked individuals, we first linked left and right flank images based on time and location and identified individuals of the wild phenotype based on their unique rosette patterns by visual comparisons, confirmed by at least two authors. We constructed year-specific spatial detection histories that indexed the capture location and daily sampling occasion of each adult individual capture. Non-independent captures (i.e. the same individual at the same location on a single sampling occasion and observations of non-independent individuals, e.g. cubs) were discarded and estimates of density pertained to adult leopards. Given that we obtained some captures of individuals whose identity was only known from a single flank (either left or right), we created two spatial detection history datasets for each year. These included captures of individuals that were identified from both flanks plus those identities that were confirmed by either flank (Both+Left and Both+Right datasets) (Augustine et al. 2018). Year-specific spatial counts of ‘unmarked’ (melanistic) individuals (*Tu*) were constructed as matrices where detectors represented rows and sampling occasions columns.

As we also wished to include capture data of unidentified sightings of marked individuals (*Tm*), we constructed year-specific spatial count datasets that incorporated captures of rosettted individuals where marks were unreadable (Figure A1) and captures where identity was only known from a single flank (either right or left). Prior to analyses, ‘capthist’ objects were constructed using package ‘secr’ in program R (R Development Core Team 2014; Efford 2020). Flank- and year-specific ‘capthist’ objects were compiled that included spatial detection history dataset (e.g. Both+Right), spatial counts of ‘unmarked’ individuals (*Tu*), the complementary spatial count dataset of unidentified sightings of marked individuals (*Tm_Left_*) and a year-specific matrix that specified individual trap layout (X and Y location in UTMs) and functionality across sampling occasions.

Population densities 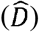 were estimated using a maximum likelihood-based SMR model implemented in package ‘secr’ within the R environment (R Development Core Team 2014; Efford & Hunter 2018; Efford 2020). We fit ‘sighting-only’ models, where the number of marked individuals was unknown, and no marking occasions were specified. As we incorporated data on unidentified sightings of marked individuals, we allowed for the estimation of a parameter for incomplete identification of marked animals when they were resighted (pID). Given that the summed counts of unmarked (*Tu*) and unidentified sightings of marked individuals (*Tm*) could result in overdispersion, we adjusted for this by first fitting the model assuming no overdispersion, then estimating the overdispersion by simulating (n = 10000) at the initial estimate and finally refitting the model using an overdispersion-adjusted pseudo-likelihood (Efford & Hunter 2018; Efford 2019). As the sampling frames across years were consistent, we created a single state-space file by applying a 15 km buffer around the trap area and generating a fine grid of equally spaced points, each representing 0.25 km^2^. We used forest cover to represent suitable and non-suitable habitat. Flank- and year-specific population densities 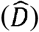, were obtained using a half-normal detection function where an intercept (g_0_) and scale parameter (sigma; σ) were estimated.

To assess the effects of pooling parameter values that could be common across years, we also implemented multi-session analyses of the data. We first merged the year-specific data for each of the flanks and created two multi-session datasets (Both+Left and Both+Right). We constructed a candidate set of eight models where g_0_, σ and pID were modelled in all parameter combinations of constant (.) and year (y) variant effects. We ranked models using Akaike’s Information Criterion adjusted for sample size (AICc; Burnham and Anderson, (2002)). All parameter inferences were based on model-averaged estimates.

Finally, we also estimated the population densities of rosetted leopards using a maximum likelihood-based SECR analysis implemented in package ‘secr’ within the R environment to compare against estimates obtained from the SMR analysis. In this analysis we excluded captures of melanistic and unidentified wild phenotype individuals (i.e. the summed counts of unmarked (*Tu*) individuals and unidentified sightings of marked individuals (*Tm*)).

## Results

From a trapping effort of 18,674 trap-days across three sampling years at MNP, we obtained 728 leopard photo-captures, of which 22.6% (165) were melanistic. While individuals were identified from both flank images in 445 (61.2%) captures, we obtained 70 (9.6%) captures of individuals identified from a single flank and 48 (6.6%) captures of unidentified rosetted individuals. The number of individuals uniquely identified from rosette patterns (M_t+1_), detections (*ndet*) used in spatial detection histories, detections of unidentified marked individuals (*Tm*) and detections of melanistic individuals (*Tu*) varied between flank and year-specific datasets (Table 1).

Leopard population density point estimates ranged from 2.61 to 4.33 individuals/100km^2^, with strong congruence between both single session and multi-session analyses (Table 2 and Figure A3). The ‘best-flank’ datasets (i.e. datasets with a higher number of individuals identified) varied between years resulting in point estimates of 4.33±SE 0.18, 2.61±0.15 and 3.37±0.14 individuals/100km^2^ for years 2016-17, 2018 and 2018-19 respectively (Table 2). Estimate precision was relatively higher for multi-session analyses (CI width 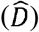; Table 2 and Figure 2). The best fit multi-session model for both flank datasets was one that incorporated constant proportion identified (pID) and year-specific intercept (g_0_) and scale parameters (σ) (Table A1). Finally, we show that point estimates of rosetted leopards obtained from an SECR analysis were 15-20% lower than estimates derived from the SMR analysis (Table A2).

## Discussion

In this study, we present an application of SMR models to estimate leopard densities when the population comprises both melanistic and wild phenotypes. The point estimates of 4.33±SE 0.18, 2.61±0.15 and 3.37±0.14 individuals/100 km^2^ across three sampling years likely represent (to our best knowledge) the first robust estimates of leopard densities from such mixed populations. Also, we show that by sharing parameters across years, our estimate precision was improved (Gerber et al. 2014).

Leopards are the most wide-ranging of large felids, yet their populations have been reduced, isolated, and even extirpated from large portions of their historic range owing to a host of threats (Jacobson et al. 2016; Rostro-García et al. 2016; Stein et al. 2020). Although the species receives research and conservation attention, there is a spatial bias in these efforts with camera trapping studies being concentrated in Africa (*P. p. pardus*) and south Asia (*P. p. fusca*) (Jacobson et al. 2016). While estimates of population density vary widely (Farhadinia et al. 2019), in the Tropical and Subtropical Moist Broadleaf Forests of Bhutan, India, Malaysia, Nepal, and Thailand, most estimates and associated detection parameters were comparable to our study (Table A3). Amongst populations that comprise both phenotypes (e.g. the Western Ghats and Eastern Himalayan moist forests; da Silva et al., (2017)), camera-trap surveys have either not estimated leopard densities (e.g. Karanth et al., (2017)), or have censored melanistic individuals from the sample, potentially biasing estimates (e.g. Borah et al. (2014); Jhala et al. (2015)). As the exclusion of melanistic captures misrepresents the population, SMR models provide an opportunity to revisit photographic capture data from past surveys towards generating estimates of leopard densities at these sites.

SMR models have been developed and implemented for a range of sampling scenarios (Sollmann et al. 2013; Efford & Hunter 2018; Whittington et al. 2018). In particular, the use of camera-trap surveys is appealing as they are cost-effective and allow for sampling carnivores that are wide-ranging, nocturnal, secretive and occur at low population densities (Alonso et al. 2015; Kane et al. 2015; Whittington et al. 2018; Jimenez et al. 2019; Murphy et al. 2019). When individuals are identified by natural markings (as in the case of leopards), it is appropriate to assume that marked individuals are a random subset of the entire population, thereby allowing for the robust estimation of population density (Royle et al. 2014; Efford & Hunter 2018). Additionally, when a greater proportion of captures are of marked individuals as opposed to unmarked animals, it is expected that the accuracy and precision of the estimates are increased (Chandler & Royle 2013). In our study, we assigned individual identity to ~70% of photo-captures, likely resulting in the high precision of our estimates. This is in contrast to studies where individuals have been identified either by artificial markings (e.g. tags, radio-collars) and/or semi-permanent natural markings (e.g. nicks, scars, physical deformities) that typically result in a greater proportion of detections of unmarked animals (Kane et al. 2015; Whittington et al. 2018). A particular shortcoming of our analysis is that we do not formally address the issues arising from partial identities in camera trapping studies (i.e., single flank photographs). While our results suggest that the variation in estimates between Both+Left and Both+Right datasets are minimal, recently developed spatial partial identity models (SPIM) could be extended to SMR models to probabilistically resolve identities and help improve estimate precision (Augustine et al. 2018, 2019).

Estimating the abundance and density of species is vital to understanding population ecology and informing species conservation. Camera-trapping in combination with SECR or SMR models provides the means to obtain such information. While IR camera traps forced into night mode in conjunction with SECR models could help estimate leopard densities when both wild and melanistic phenotypes exist, SMR models allow for the estimation of density with photo capture data obtained from camera trap surveys conducted using only white-flash units. Additionally, SMR models enable researchers to revisit historic camera trap survey data, thereby generating valuable information on populations. Finally, these models can also be applied to estimate the population densities of other cryptic carnivores that exhibit similar phenotypic polymorphism such as Jaguars *Panthera onca* and Oncilla *Leopardus tigrinus* (Silveira et al. 2010; Mooring et al. 2020).

## Acknowledgements

We are thankful to the Forest Department, Government of Assam and Bodoland Territorial Council for permitting us to carry out the survey, and Assam University, Diphu Campus for academic support to the second author. We are grateful to Mr Anindya Swargowari, IFS, Additional Principal Chief Conservator of Forests cum Council Head of the Department of Forest, BTC, Mr Hiranya Kumar Sarma, IFS, former Field Director, Mr Amal Chandra Sarmah, IFS, current Field Director, R. N Boro, deputy Director, IFS, Abbas Dewan, ACF, of the Manas Tiger Reserve, the Range officers of the MNP, Babul Brahma, Kunja Basumatary, Pranab Das, Kameswar Baro and the frontline staff for providing logistic support during the field surveys. At Aaranyak, we thank Dr B. K. Talukdar, CEO & SG and Dr B. P. Lahkar for their continued support. We thank Dr Anupam Sarmah, Head, Brahmaputra Landscape and WWF network for their support. We are thankful to D.D. Boro, Kiran Ch. Basumatary, for sharing field knowledge and administrative support. This study was made possible through field support by Arif Hussain, Dr Navaneethan Balasubramaniam, Abinash Parida, Nanka Lakra, Sansuma Narzari, Hiranya Moran, Utpal Das, Nandeswar Wary, Pabitra Sutradhar, Amal Pathak, Bhaskor Barukial, Binita Baruwati, Apurba Das, Debasish Buragohain, Prosenjit Sheel, Pronit Basumatary, Mukesh Kherkatari, Ratul Das, Mujamil Haque, Bipul Ch. Nath, Ranjit Urang and Dilli Boro. Aaranyak is thankful to Integrated Tiger and Habitat Conservation Programme of IUCN–KfW, Panthera, RTCF Grant of the US Fish and Wildlife Service and WWF for financial support to carry out this study. Chris Hallam, Hugh Robinson, Bart Harmsen, Matthew Rogan, Ben Augustine, Mousumi Ghosh-Harihar, Dan Parker and two anonymous reviewers are thanked for their comments and suggestions on previous drafts of the manuscript.

---

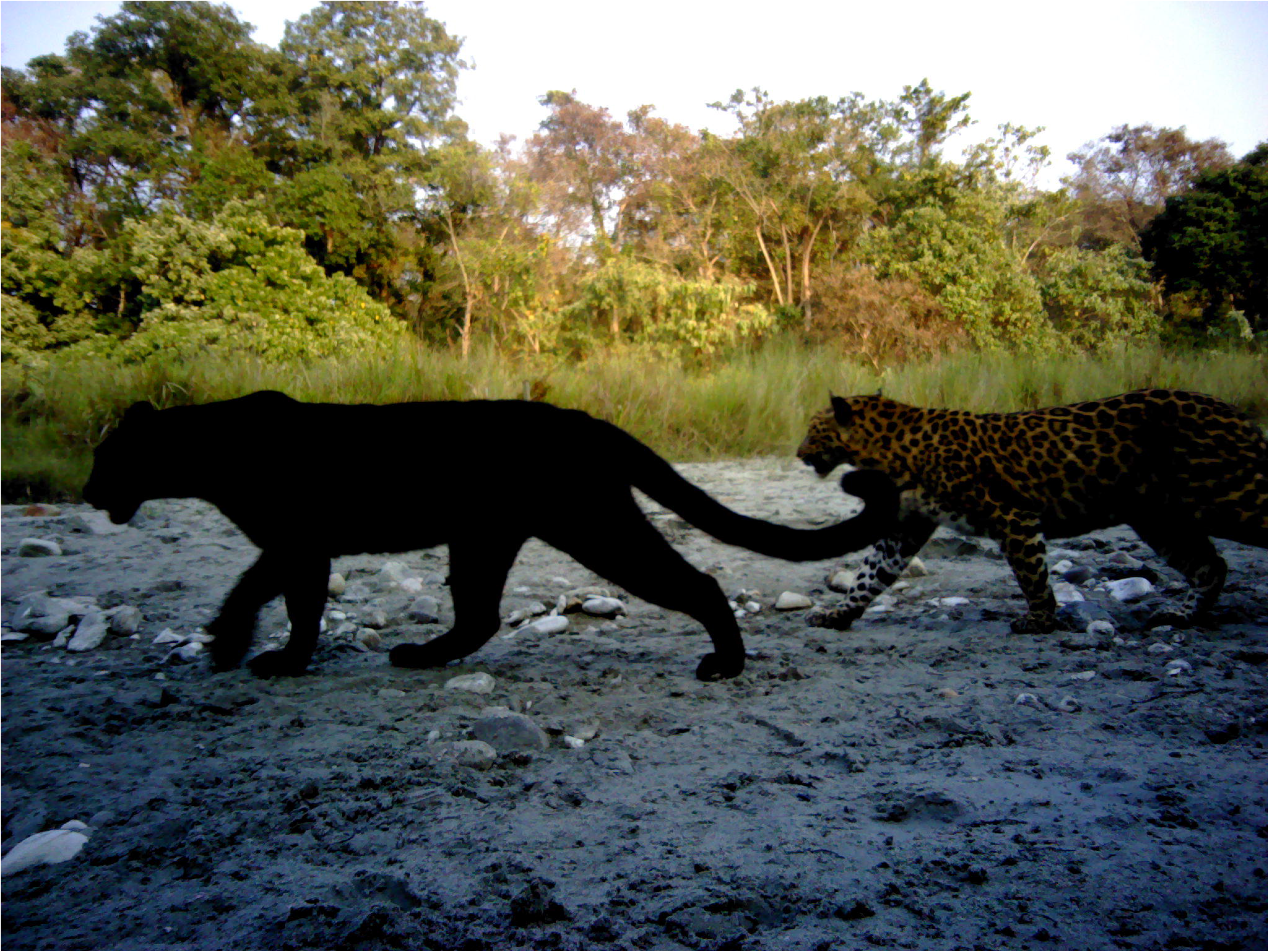

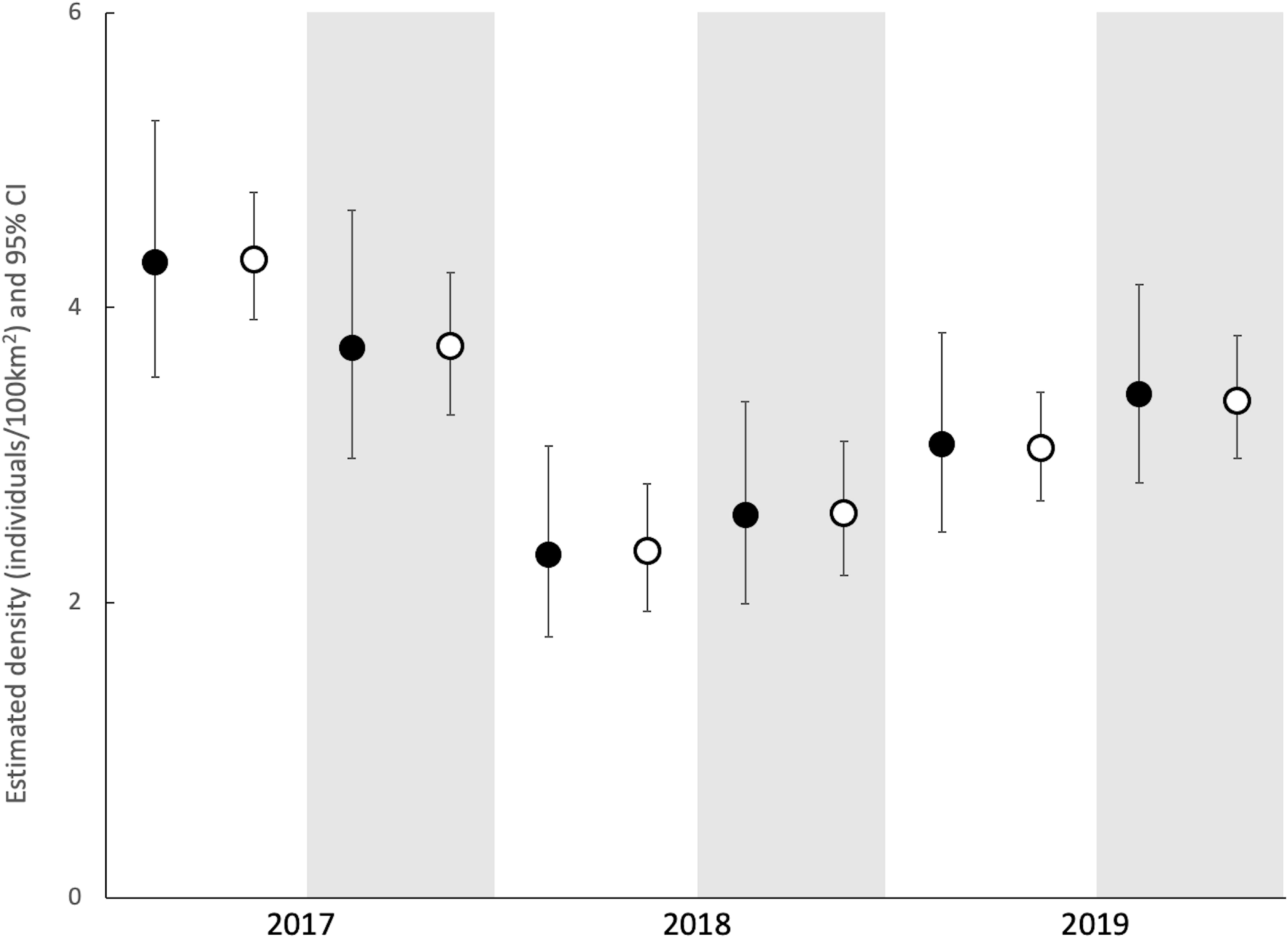

