## Appendix for "Population density modeling of mixed polymorphic phenotypes: an application of spatial mark-resight models"

**Figure A1. Example images of unidentified marked individuals.**

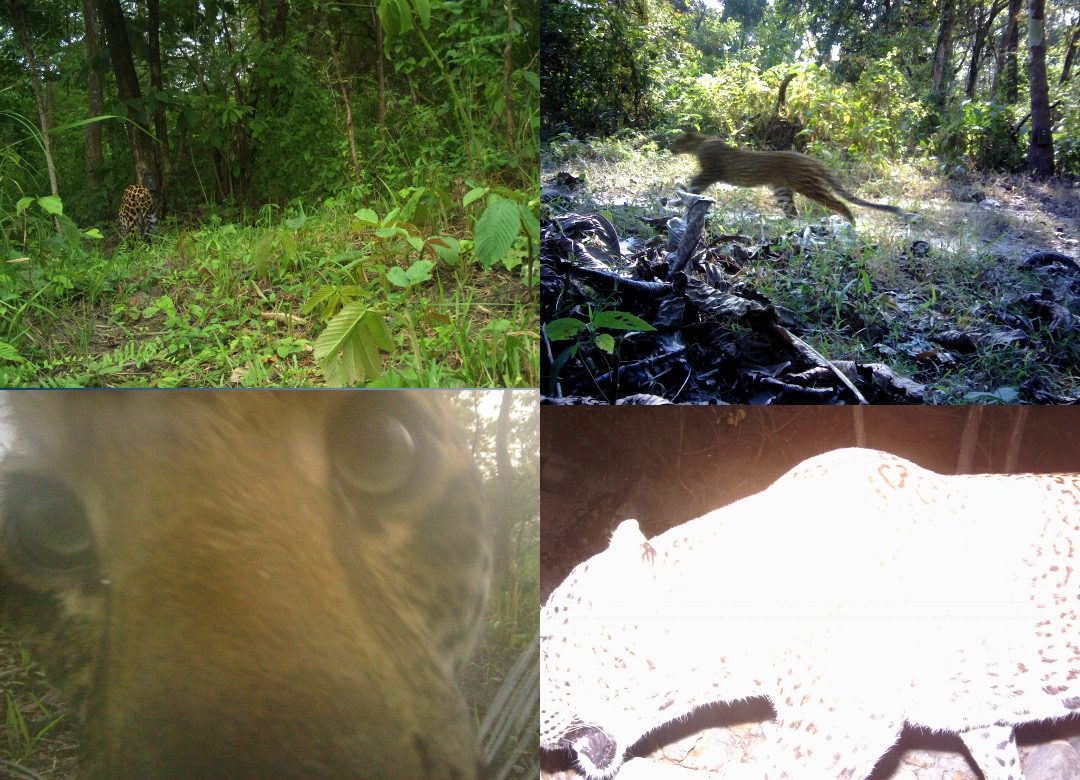

**Figure A2. Maps of Manas National Park indicating camera trap locations for the three sampling years.**

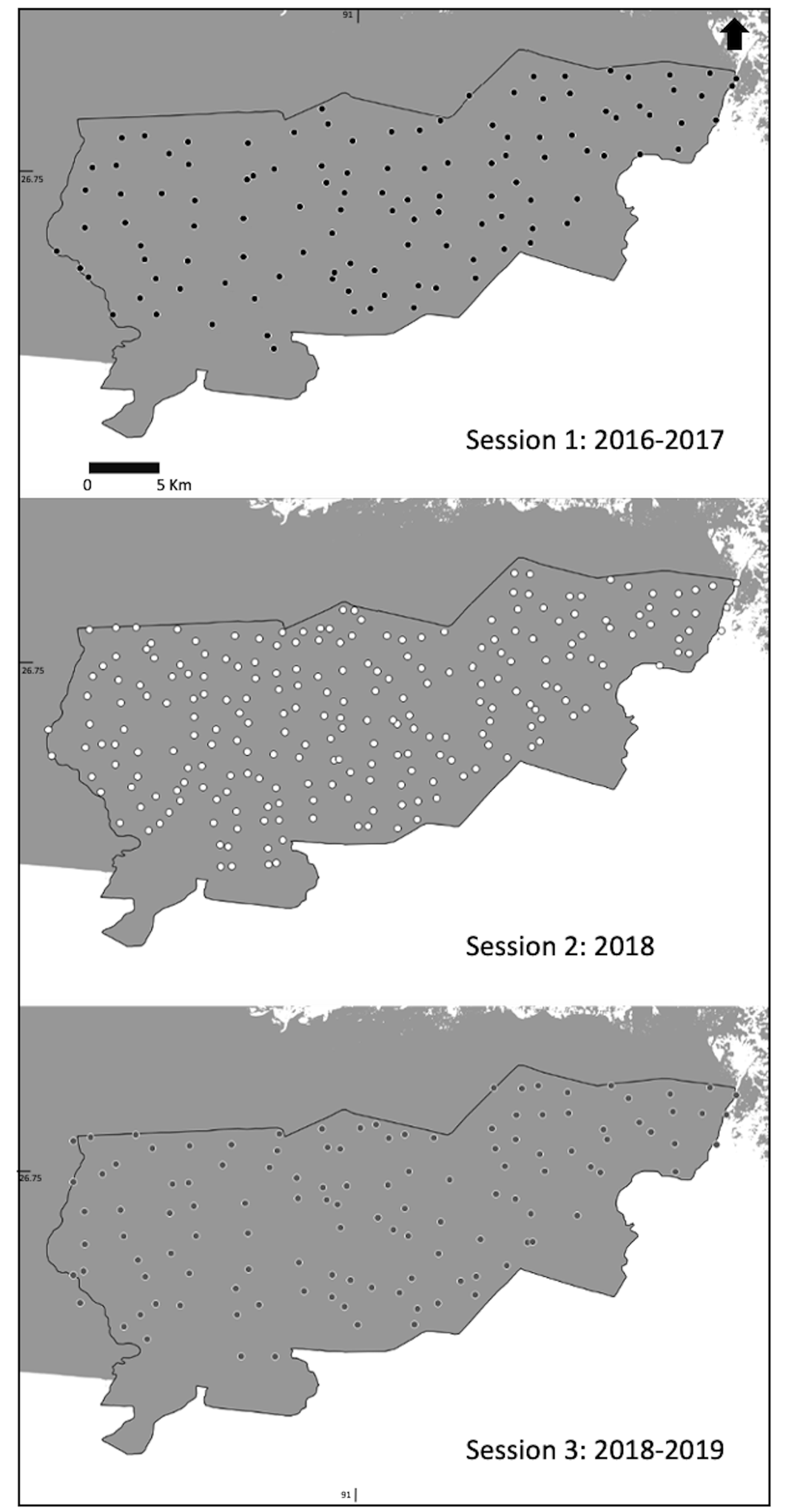

**Table A1. Model selection table for the multisession analyses of leopard data using Spatial Mark Resight models over three years at Manas National Park.** Presented are the results for the analyses of Both + Left and Both + Right multisession datasets. Models are ranked in order of AICc; Akaike’s Information Criterion adjusted for sample size with the best model in the first row (lowest AICc) for Both + Left analyses. Presented are the ΔAICc (difference in AICc from the best model), AICc weights and number of parameters. The model combinations include constant (.) and year (y) variant effects on the intercept (g0), scale (sigma) and the proportion identified (pID) parameters.

| Model | No. of parameters | Both + Left | | |  | Both + Right | | |
| --- | --- | --- | --- | --- | --- | --- | --- | --- |
|  |  | AICc | ΔAICc | AICc weight |  | AICc | ΔAICc | AICc weight |
| g0 (y) sigma (y) pID (.) | 10 | 8688.592 | 0 | 0.4354 |  | 8661.594 | 0 | 0.6522 |
| g0 (y) sigma (y) pID (y) | 12 | 8689.348 | 0.756 | 0.2983 |  | 8667.068 | 5.474 | 0.0422 |
| g0 (y) sigma (.) pID (.) | 8 | 8691.958 | 3.366 | 0.0809 |  | 8663.901 | 2.307 | 0.2058 |
| g0 (y) sigma (.) pID (y) | 10 | 8692.308 | 3.716 | 0.0679 |  | 8669.094 | 7.5 | 0.0153 |
| g0 (.) sigma (.) pID (y) | 8 | 8692.691 | 4.099 | 0.0561 |  | 8670.779 | 9.185 | 0.0066 |
| g0 (.) sigma (.) pID (.) | 6 | 8693.006 | 4.414 | 0.0479 |  | 8666.244 | 4.65 | 0.0638 |
| g0 (.) sigma (y) pID (y) | 10 | 8696.905 | 8.313 | 0.0068 |  | 8673.974 | 12.38 | 0 |
| g0 (.) sigma (y) pID (.) | 8 | 8696.945 | 8.353 | 0.0067 |  | 8669.277 | 7.683 | 0.014 |

**Figure A3. Spatial Mark Resight model parameter estimates for the three sampling years.** Estimates of the intercept (g0), scale (sigma) and proportion identified (pID) parameters for the single session analyses (closed circles; ⚫) and multisession analyses (open circles; ⭘) for the Both + Left and Both + Right datasets. Error bars indicate 95% confidence intervals around estimates.

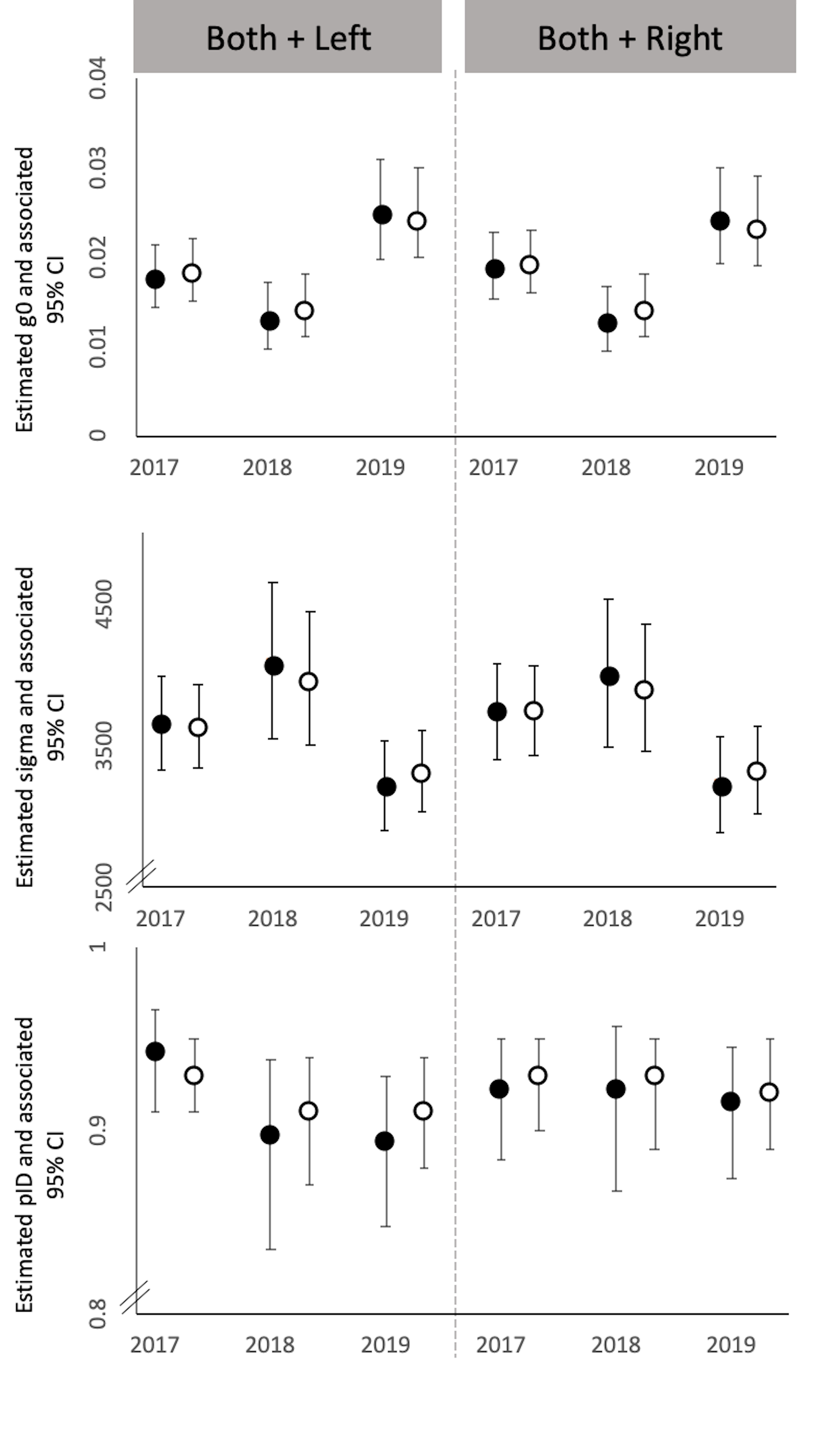

**Table A2.** **Spatially Explicit Capture-Recapture (SECR) estimates of rosetted leopards over three years at Manas National Park.** Parameter estimates from single session SECR analyses of rosetted leopard data over three sessions and for each of the flank datasets. Presented are the estimated density ($\hat{D}$; individuals/100km^2^), intercept parameter ($\hat{g0}$), scale parameter in meters ($\hat{\sigma}$) with associated standard errors (SE) and the Goodness-of-fit test statistic.

| Years | Flank dataset | D (SE) | g0 (SE) | sigma (SE) | GOF |
| --- | --- | --- | --- | --- | --- |
| 2016-17 | Both + Left | 3.51 (0.63) | 0.017 (0.002) | 3609.8 (161.7) | 0.955 |
|  | Both + Right | 3.01 (0.57) | 0.018 (0.002) | 3725.1 (173.3) | 0.801 |
| 2018 | Both + Left | 1.89 (0.1) | 0.012 (0.002) | 4050.4 (276.3) | 0.689 |
|  | Both + Right | 2.13 (0.1) | 0.012 (0.002) | 3962.8 (261.6) | 0.718 |
| 2018-19 | Both + Left | 2.62 (0.09) | 0.023 (0.003) | 3181.7 (155.9) | 0.964 |
|  | Both + Right | 2.94 (0.1) | 0.023 (0.003) | 3189.5 (168.3) | 0.997 |

**Table A3. Comparison of leopard density estimates from Tropical and Subtropical Moist Broadleaf Forests of south and south-east Asia.** Presented are the site name, country, estimated population density and citation to the study.

| S. No. | Site | Country | Density  (individual/100 km2) | Sigma (SE/SD/95% credible intervals)  meters | g0/lambda0 (SE/SD/95% credible intervals) | Reference |
| --- | --- | --- | --- | --- | --- | --- |
| 1 | Jigme Singye Wangchuck National Park^a,^ * | Bhutan | 1.04 (0.01) | Not applicable | | (Wang & Macdonald 2009) |
| 2 | Royal Manas National Park^b,^ * | Bhutan | 10 | 2200 (1560, 3100) M;  1200 (960, 1550) F | 0.03 (0.016, 0.047) | (Goldberg et al. 2015) |
| 3 | Manas National Park* | India | 3.4(0.82) | 1.602 (0.2) | 0.86 (0.34) | (Borah et al. 2014) |
| 4 | Manas National Park* | India | 4.33 (0.18) | 3627.07 (150.69) | 0.018 (0.002) | This study^c^ |
| 5 | Manas National Park* | India | 2.61 (0.15) | 3886.35 (228.33) | 0.014 (0.002) |  |
| 6 | Manas National Park* | India | 3.37(0.14) | 3319.48 (157.45) | 0.023 (0.003) |  |
| 7 | Pakke Tiger Reserve* | India | 2.82(1.2) | Not presented | | (Selvan et al. 2014) |
| 8 | Kalakad-Mudanthurai Tiger Reserve* | India | 1.3 (1.0) | 2800 (1200) | 0.022 (0.014) | (Ramesh et al. 2012) |
| 9 | Kalakad-Mudanthurai Tiger Reserve* | India | 2.8 (2.0) | 1500 (470) | 0.014 (0.008) |  |
| 10 | Parsa Wildlife Reserve^b^ | Nepal | 3.78 (0.85) | 2880 (400) | 0.02 (0.005) | (Thapa et al. 2014) |
| 11 | Kenyir Wildlife Corridor* | Malaysia | 3.31 (1.14) | 2249 M;  2699 F | 0.014 M;  0.0039 F | (Hedges et al. 2015) |
| 12 | Thung Yai Naresuan (East) Wildlife sanctuary* | Thailand | 0.66 (0.27) | 6490 (1411) | 0.0090 (0.004) | (Vinitpornsawan 2013) |

*- Sites that hare known to have melanistic phenotypes

^a^ – Estimate based on a non-spatial capture recapture analysis (therefore

^b^ – Inference based on a Bayesian analysis of spatial capture recapture data

^c^ – Estimates derived from Spatial Mark-Resight models incorporating melanistic captures.
